# Sperm-contributed centrioles segregate stochastically into blastomeres of 4-cell stage *C. elegans* embryos

**DOI:** 10.1101/2023.01.27.525957

**Authors:** Pierre Gönczy, Fernando R. Balestra

## Abstract

Whereas both sperm and egg similarly contribute nuclear genetic material to the zygote of metazoan organisms, the inheritance of other cellular constituents is unequal between the two gametes. Thus, the centriole organelle is contributed solely by the sperm to the zygote in most systems. Whereas centrioles can have a stereotyped distribution during some asymmetric divisions, whether sperm-contributed centrioles are distributed in a stereotyped manner in the resulting embryo is not known. Here, we addressed this question in *Caenorhabditis elegans* using marked mating experiments, whereby the presence of sperm-contributed centrioles is monitored in the resulting embryos using the stable centriolar component SAS-4::GFP, as well as GFP::SAS-7. Our analysis demonstrates that the distribution of sperm-contributed centrioles is stochastic in 4-cell stage embryos. Moreover, using sperm from *zyg-1* mutant males that harbor a single centriole, we show that the older sperm-contributed centriole is also distributed stochastically in the resulting embryo. Overall, we conclude that in contrast to the situation during some asymmetric cell divisions, centrioles contributed by the male germ line are distributed stochastically in embryos of *C. elegans*.

## Introduction

Whereas the genetic material is contributed in an equal manner to the zygote by the two parental gametes in metazoan organisms, this is not the case for the remainder of the cellular material. In most species, the egg contributes the bulk of the mRNAs, proteins, and cytoplasmic constituents to the zygote. The same holds true for mitochondria, which are inherited and retained strictly maternally by the developing embryo. Conversely, the sperm is the sole contributor of centrioles, which are absent from the egg. Such differential contribution of centrioles by the two gametes is critical for endowing the zygote with the correct number of centrioles at the onset of life. Whether sperm-contributed centrioles are segregated thereafter to specific cells of developing embryos is not known.

Centrioles are small microtubule-based organelles that are critical for fundamental cellular processes across eukaryotes (reviewed by Bornens 2012; Winey and O’Toole 2014). Centrioles are key notably in their capacity as basal bodies that seed the formation of cilia and flagella in specific cell types, including sperm cells in most species. Moreover, in animal systems, centrioles recruit the pericentriolar material (PCM), thus forming the centrosome, which acts as an important microtubule organizing center (MTOC). Through this role, centrioles are important for core cellular processes such as polarity and division.

Centrosomes have an inherent cell generational asymmetry: most proliferating cells are born with two centrioles, a so-called mother centriole that has been generated at least one cell generation prior, and a so-called daughter centriole, which has been generated during the previous cell cycle (reviewed by Nigg and Stearns 2011). Towards the onset of S phase, both mother and daughter centriole seed the formation of a procentriole in their vicinity, yielding two centrosomes, each containing a centriole/procentriole pair, which direct bipolar spindle assembly during mitosis.

Interestingly, centriole inheritance is stereotyped in some cases of asymmetric cell division (reviewed by Yamashita 2009). For instance, in germ line stem cells of the *Drosophila* testis, the centrosome harboring the older centriole, which is referred to as the older centrosome, is inherited systematically by the cell that retains the stem cell fate (Yamashita *et al.* 2007). Perturbing such stereotyped inheritance through mutation of the PCM protein centrosomin randomizes centrosome distribution and results in declined stem cell function (Yamashita *et al.* 2007). Conversely, the older centrosome is inherited systematically by the ganglion mother cell during asymmetric division of *Drosophila* neuroblasts, whereas stem-like neuroblasts retain the younger centrosome (Conduit and Raff 2010; Januschke *et al.* 2011). Such stereotyped asymmetric centriole inheritance is not limited to *Drosophila.* Thus, radial glial progenitors in the ventricular zone of the developing mouse cortex also exhibit stereotyped centrosome inheritance (Wang *et al.* 2009). In this case, the mother centriole is inherited systematically by the radial glial progenitor stem cell during asymmetric division, reminiscent of the situation in the fly testis. Moreover, depletion of the mother centriole-specific protein Ninein, which is essential for mother centriole anchoring to the plasma membrane, randomizes centrosome distribution in this setting. Importantly, this is accompanied by premature depletion of radial glial progenitor from the ventricular zone (Wang *et al.* 2009).

In contrast to the wealth of information regarding the inheritance pattern of centrosomes and its importance in stem cell systems, little is known regarding the distribution of the two sperm-contributed centrioles in the resulting developing embryos. Conceivably, such inheritance could be stereotyped as well and thereby potentially endow specific blastomeres with paternally-derived components. This paucity of information stems notably from the fact that centrioles assembled in the zygote using maternal components can be difficult to distinguish from those contributed by sperm. This experimental limitation can be circumvented with so-called marked mating experiments, which have been deployed in *Caenorhabditis elegans* (Kirkham *et al.* 2003; Leidel and Gönczy 2003; Balestra *et al.* 2015). In this setting, a stable centriolar component is labelled in the sperm with a fluorescent protein such as GFP. Upon mating with an egg devoid of this GFP-tagged centriolar component, the two sperm-contributed centrioles stably harbor GFP, whereas all centrioles generated in the zygote do not. Therefore, this experimental setting offers a unique means to monitor the fate of paternally contributed centrioles in early embryos.

## Results

### Experimental design

We set out to use marked mating experiments to test whether sperm-contributed centrioles are distributed in early *C. elegans* embryos in a stereotyped fashion, or instead stochastically, focusing our analysis on the 4-cell stage. The experimental strategy is summarized schematically in Figure 1. Hermaphrodites expressing TagRFP-T::SAS-7 and homozygous for *fem-1(hc17ts)* are raised at 25°C, which results in them lacking sperm. Such feminized hermaphrodites are then mated to males expressing endogenously tagged SAS-4::GFP or GFP::SAS-7 (Fig. 1A). As a result, the two sperm-contributed centrioles are marked with GFP (Fig. 1B), and each such centriole seeds the formation of one procentriole in its vicinity during the first cell cycle in the zygote using maternal-contributing components given that zygotic transcription begins only later. The two resulting centrosomes, each with a centriole/procentriole pair, constitute the two poles of the mitotic spindle at the end of the first cell cycle (Fig. 1C, left). Each resulting daughter cell, termed AB and P_1_, respectively, will necessarily inherit one of the two paternally contributed centrioles (Fig. 1C, right). At the onset of the second cell cycle, each of the four centrioles that are present at that time then seeds the formation of one procentriole in its vicinity (Fig. 1C, right). As illustrated in Figure 1D, this could in principle yield four distinct distributions of sperm-contributed centriole at the following cell cycle, in 4-cell stage embryos: ABa and P_2_, AB and EMS, ABp and P_2_, or ABp and EMS.

**Figure 1.**
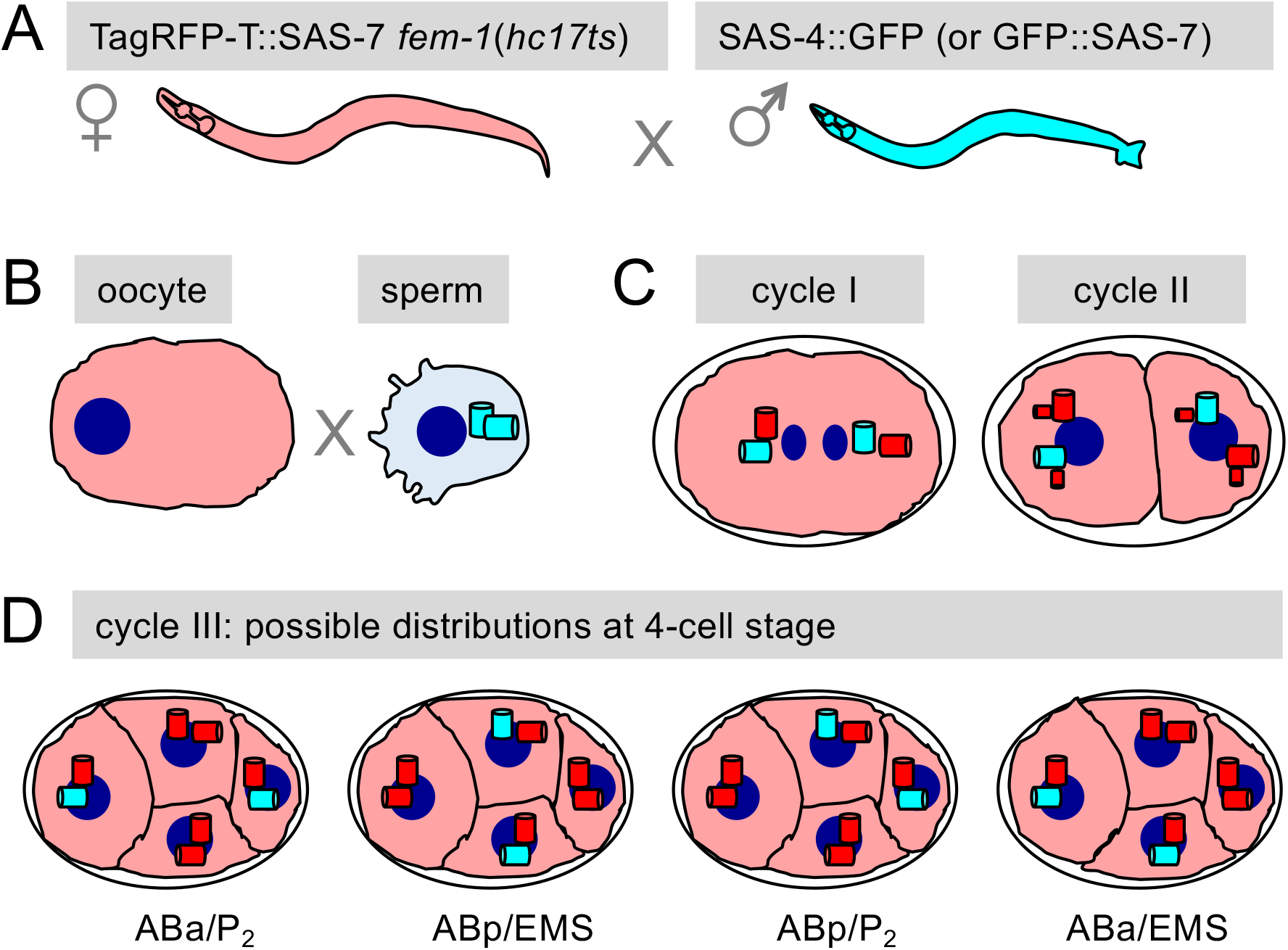
Experimental design. **(A, B)** Hermaphrodites expressing TagRFP-T::SAS-7 (represented in red) and homozygous for *fem-1*(*hc17ts*) raised at 25°C lack sperm. Such feminized hermaphrodites are mated to males expressing SAS-4::GFP or GFP::SAS-7 (both represented in cyan), and the fate of GFP-marked sperm-contributed centrioles analyzed in the resulting embryos. **(C)** At the end of the first cell cycle (left), a bipolar spindle assembles, with each spindle pole harboring one sperm-contributed centriole (cyan) and one procentriole assembled in the zygote with maternally contributed components (red). At the onset of the second cell cycle (right), each of the four centrioles seeds the formation of one procentriole in its vicinity (red). **(D)** Possible distributions of sperm-contributed centrioles in the third cell cycle (4-cell stage embryos): ABa and P_2_, AB and EMS, ABp and P_2_, or ABp and EMS. For simplicity, procentrioles forming in the vicinity of each centriole are not represented here.

### Stochastic distribution of sperm-contributed centrioles at the 4-cell stage

We deployed the above experimental strategy using primarily SAS-4::GFP because prior work established that this evolutionarily conserved centriolar protein does not undergo significant exchange once incorporated in the centriole (Kirkham *et al.* 2003; Leidel and Gönczy 2003; Balestra *et al.* 2015). As illustrated in Figure 2A and 2B, as well as in Figure S1A, the TagRFP-T::SAS-7 signal serves to identify the location of centrioles in each cell, whereas the SAS-4::GFP signal is monitored to determine the distribution of sperm-derived centrioles. Importantly, this analysis uncovered that the fraction of 4-cell embryos harboring a sperm-contributed centriole in ABa is indistinguishable from that harboring a sperm-contributed centriole in ABp (Fig. 2B-F; see Table S1 for statistical analysis; n=257 embryos). Similarly, the fraction of 4-cell embryos with a sperm-contributed centriole in EMS is indistinguishable from that with a sperm-contributed centriole in P_2_ (Fig. 2B-F; Table S1). Moreover, there is no stereotyped pair-wise arrangement of cells harboring sperm-contributed centrioles; instead, the fraction of embryos with sperm-contributed centrioles in ABa and EMS is indistinguishable from that in ABp and EMS, ABa and P_2_, or ABp and P_2_ (Fig. 2G; Table S1).

**Figure 2.**
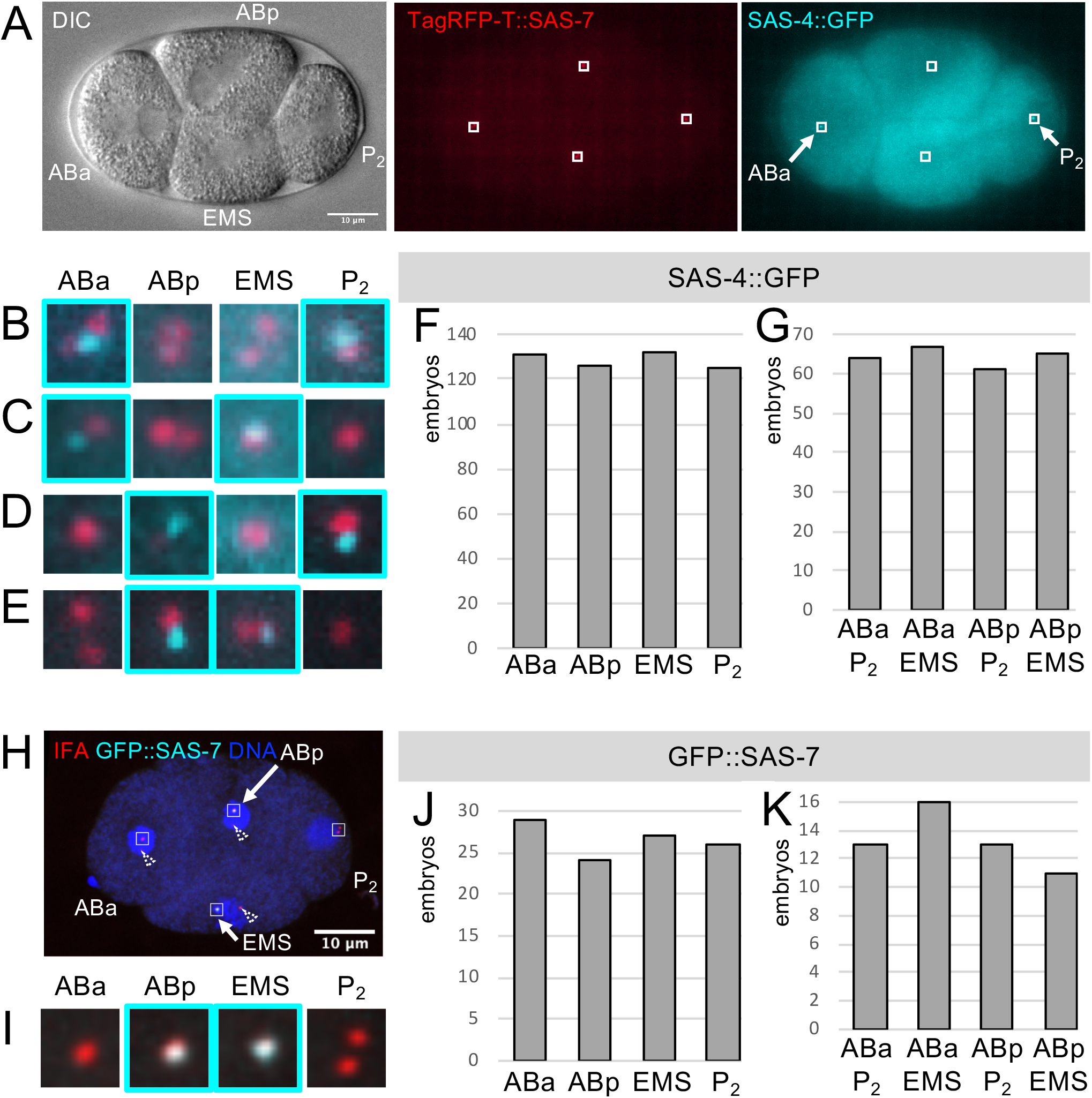
Stochastic distribution of sperm-contributed centrioles in 4-cell stage embryos. **(A)** DIC (left), TagRFP-T::SAS-7 (center, red) and SAS-4::GFP (right, cyan) live imaging of 4-cell stage embryo resulting from mating of TagRFP-T::SAS-7 *fem-1*(*hc17ts*) hermaphrodites with males expressing SAS-4::GFP. The ABa, ABp, EMS and P_2_ blastomeres are indicated in the DIC image. Small boxes indicate insets magnified in B. Arrows point to the ABa and P_2_ blastomeres, which inherited the two sperm-contributed centrioles in this particular embryo. **(B)** Dual color (TagRFP-T::SAS-7, red; SAS-4::GFP, cyan) magnified insets of four regions indicated in A; note that SAS-4::GFP is present in ABa and P_2_, as emphasized by the dashed cyan contours. **(C-E)** Analogous insets from embryos shown in Fig. S1B-D, with SAS-4::GFP present in ABa and EMS (C), ABp and P_2_ (D) or ABp and EMS (E). **(F, G)** Distribution of sperm-contributed centrioles marked by SAS-4::GFP in 4-cell stage embryos. N=257 embryos, 219 analyzed by live imaging, 32 following ethanol fixation, 6 after immunofluorescence analysis. Note that 11/257 embryos were fertilized by *zyg-1*(*b1ts*) sperm grown at the permissive temperature, hence also contributing two SAS-4::GFP-marked centrioles. Overall occurrences (F): ABa −131, ABp −126, EMS −132, P_2_ −125. Overall occurrences (G): ABa and P_2_ −64; ABa and EMS −67; ABp and P_2_ −61; ABp and EMS −65. See Table S1 for statistical analysis. **(H)** Confocal microscopy of 4-cell stage embryo resulting from mating of TagRFP-T::SAS-7 *fem-1*(*hc17ts*) hermaphrodites with males expressing GFP::SAS-7, stained with antibodies against the centriolar marker IFA (red) and GFP (cyan); DNA counterstain is in blue. Small boxes indicate insets magnified in I. Arrows point to the ABp and EMS blastomeres, which inherited the two sperm-contributed centrioles in this particular embryo. Dashed arrowheads point to centrioles outside the magnified areas, some of which are poorly visible in the displayed optical slices. **(J, K)** Distribution of sperm-contributed centrioles marked by GFP::SAS-7 in 4-cell stage embryos. N=53 embryos, 3 analyzed by live imaging, 14 following ethanol fixation, 36 after immunofluorescence analysis. Overall occurrences (J): ABa −29; ABp −24; EMS −27; P_2_ −26. Overall occurrences (K): ABa and P_2_ −13; ABa and EMS −16; ABp and P_2_ −13; ABp and EMS −11. See Table S1 for statistical analysis.

To test whether this outcome is specific to SAS-4::GFP, we conducted analogous experiments using GFP::SAS-7, which we found to exhibit a strong signal in sperm centrioles, making it a potentially suitable candidate for marked mating experiments. We found, however, that part of the GFP::SAS-7 signal present in sperm centrioles was lost shortly after fertilization, such that we relied primarily on immunofluorescence analysis to more reliably detect sperm-contributed centriolar GFP::SAS-7 in 4-cell stage embryos. We used antibodies against the centriolar marker IFA to identify all centrioles, as well as against GFP to detect specifically sperm-contributed centrioles (Fig. 2H, 2I). This analysis demonstrated that, just like for SAS-4::GFP, the distribution of sperm-contributed centrioles is indistinguishable when comparing ABa with ABp, as well as EMS with P_2_ (Fig. 2J; Table S1; n=53 embryos). Moreover, no stereotyped pair-wise arrangement is observed in this case either (Fig. 2K; Table S1). Taken together, these experiments lead us to conclude that paternally contributed centriole are distributed in a stochastic fashion in 4-cell stage *C. elegans* embryos.

### The older sperm-contributed centriole can be present in any blastomere at the 4-cell stage

We next set out to address whether there might be a preferential inheritance of the older versus the younger sperm-contributed centriole in the resulting embryos. However, no marker is known to distinguish the older centriole from the younger one in *C. elegans.* Proteins such as Ninein that are present specifically at the mother centriole in other systems are absent from the worm genome, and the two sperm-contributed centrioles appear similar by electron-microscopy (EM) (Wolf *et al.* 1978). To bypass this limitation, we designed a genetic strategy relying on the temperature-sensitive mutant allele *zyg-1*(*b1ts*) (Wood *et al.* 1980). ZYG-1 is a kinase essential for procentriole formation in *C. elegans* and a relative of the Polo-like-kinase PLK4 that exerts a similar function in other metazoan organisms (O’Connell *et al.* 2001). The *zyg-1*(*b1ts*) mutant allele is endowed with normal function at the permissive temperature of 15°C, but results in a strong and perhaps complete loss of function phenotype upon shifting to the restrictive temperature of 25°C (Wood *et al.* 1980; O’Connell *et al.* 2001).

In this modified marked mating experiments, we shifted *zyg-1*(*b1ts*) mutant males from 15°C to 25°C during the third larval instar stage. This enables assembly of procentrioles in the proliferating mitotic zone of the gonad prior to the temperature shift, but prevents their formation thereafter. As a result, most sperm cells derived from such animals contain a single centriole, which correspond to the older one in the wild-type, whereas the centriole that should have been generated next is missing (O’Connell *et al.* 2001). During the first cell cycle, this single sperm-contributed centriole seeds the formation of a procentriole in its vicinity. Because this centriole/procentriole pair remains together until the end of the first cell cycle, a monopolar spindle assembles, leading to aberrant chromosome segregation and cleavage failure (Fig. 3A, left-most). In the second cell cycle that follows, each centriole seeds the formation of a new procentriole, leading to bipolar spindle assembly (Fig. 3A, second panel), and the generation of AB-like and P_1_-like cells in the third cell cycle (Fig. 3A, third panel). These two blastomeres then divide asynchronously, with the AB-like cell undergoing mitosis before the P_1_-like cell, as is the case for AB and P_1_ in the second cell cycle of wild-type embryos, ultimately yielding embryos that harbor AB-like, ABp-like, EMS-like and P_2_-like blastomeres (Fig. 3A, right-most).

**Figure 3.**
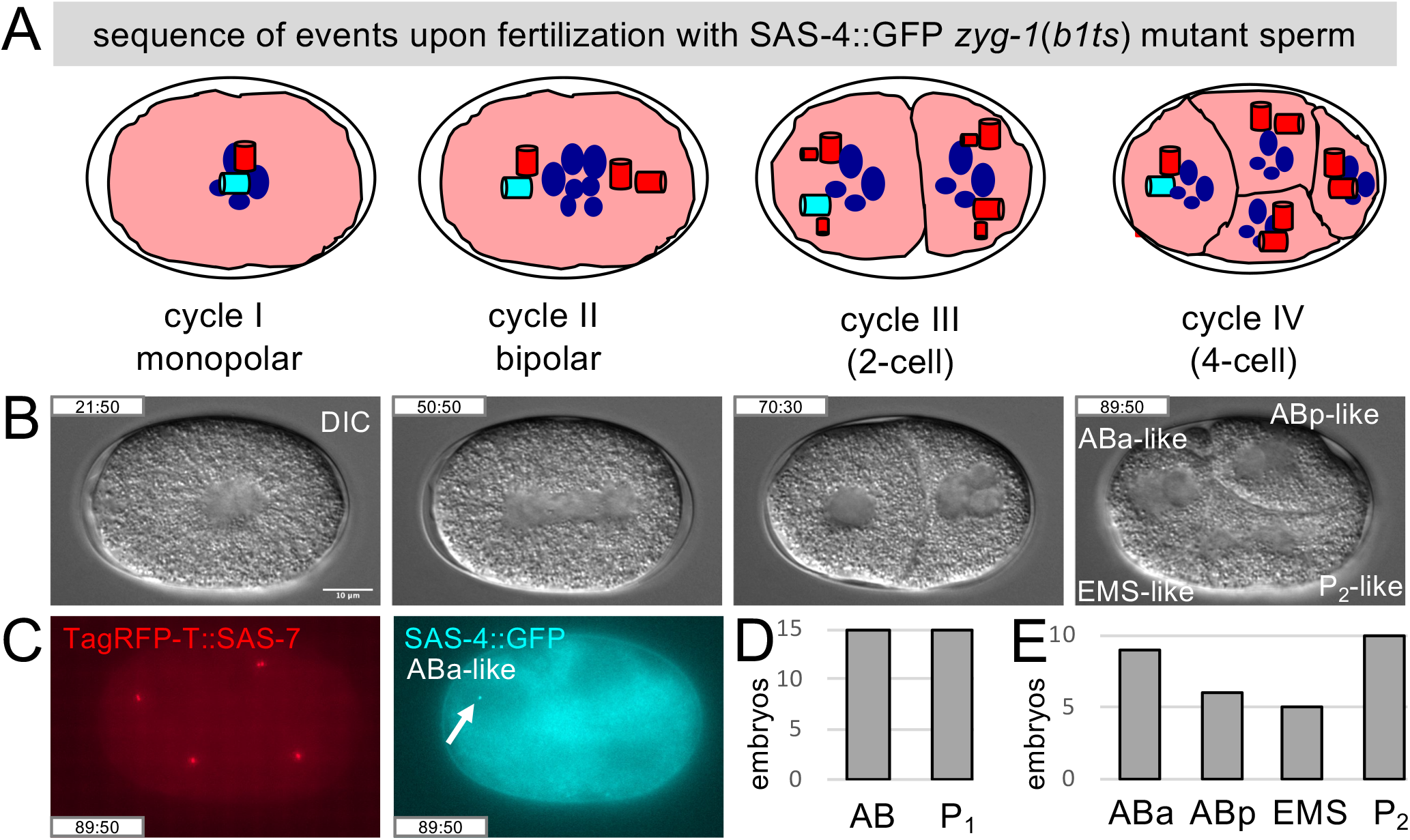
The older sperm-contributed centriole can be present in any blastomere at the 4-cell stage. **(A)** Schematic of sequence of events in embryos following fertilization of hermaphrodites expressing TagRFP-T::SAS-7 and homozygous for *fem-1*(*hc17ts*), raised at 25°C and thus lacking sperm, by *zyg-1*(*b1ts*) mutant males shifted to 25°C during spermatogenesis, leading to sperm with only one centriole marked by SAS-4::GFP. During the first cell cycle, this single centriole seeds the formation of a procentriole in its vicinity, but the first division is monopolar, leading to aberrant chromosome segregation and cleavage failure (left). In the second cell cycle, each centriole seeds the formation of a procentriole, such that a bipolar spindle assembles during mitosis (second panel), generating AB-like and P_1_-like cells in the third cell cycle (third panel). These cells divide asynchronously, with the AB-like cell going first, yielding embryos at the fourth cell cycle that harbor AB-like, ABp-like, EMS-like and P_2_-like blastomeres (right). Color code as in Fig. 1A. **(B)** DIC images from a time-lapse recording corresponding to the stages schematized in (A), with an indication of the resulting AB-like, ABp-like, EMS-like and P_2_-like blastomeres (at time 89:50; time in min:sec since the beginning of the recording). See also corresponding Video S1. **(C)** TagRFP-T::SAS-7 (left, red) and SAS-4::GFP (right, cyan) live imaging of embryo shown in (B) (at time 89:50). The arrow points to the sole sperm-contributed centriole, which is present in the ABa-like blastomere in this particular embryo. **(D, E)** Corresponding distribution in 4-cell embryos of sole sperm-contributed centriole marked by SAS-4::GFP; panel D reports the sum of sperm-contributed centrioles in AB-derived and P_1_-derived blastomeres, panel E their presence in AB-like, ABp-like, EMS-like and P_2_-like blastomeres. Overall occurrences: ABa-like −9; ABp-like −6; EMS-like −5; P_2_-like −10. See Table S1 for statistical analysis.

Using live imaging of the entire sequence of events to ensure proper staging of embryos (Fig. 3B, 3C; Video S1), we addressed whether the single centriole contributed by sperm from *zyg-1*(*b1ts*) mutant males and harboring SAS-4::GFP segregates strictly to AB-derived or P_1_-derived cells, scoring blastomeres at the 4-cell stage. As shown in Figure 3D and 3E, we found this not to be the case, as this single marked centriole can be present in either ABa-like, ABp-like, EMS-like or P_2_-like blastomere (Table S1). We conclude that the older sperm-contributed centriole is also distributed in a stochastic fashion in 4-cell stage embryos.

## Discussion

Our findings demonstrate that, in contrast to the stereotypy of centriole distribution observed during some asymmetric divisions in flies and mammals, sperm-contributed centrioles are distributed in a stochastic manner in embryos of *C. elegans.* Therefore, although sperm-contributed centrioles persist for several hours during *C. elegans* embryogenesis (Balestra et al. 2015), the present findings estatblish that they cannot endow specific blastomeres with potential trans-generational biological information. Interestingly, experiments with Dendra2::SAS-4 showed that the older centriole marked following photo-conversion can also segregate to both daughter cells in the progeny of ABprpppaa and ABprpppap in *C. elegans* (Erpf and Mikeladze-Dvali 2020), indicating that stochastic inheritance in the worm is not restricted to sperm-contributed centrioles.

It will be interesting to investigate whether the random distribution of sperm-contributed centrioles uncovered here is a species-specific feature, especially given that nematode sperm is not flagellated, in contrast to the situation in most other metazoan species. Moreover, whereas the two *C. elegans* sperm-contributed centrioles are analogous at the ultrastructural level (Wolf *et al.* 1978), they are different from one another in other species, including *Drosophila* and man (Fawcett 1975; Blachon *et al.* 2009). It is plausible that in those cases such ultrastructural differences translate into stereotyped inheritance of sperm-contributed centrioles in the resulting embryos. Alternatively, the stochastic distribution revealed in our study might reflect a more general naïve organelle state characteristic of sperm cells prior to fertilization.

## Figure and Video legends

**Figure S1.**
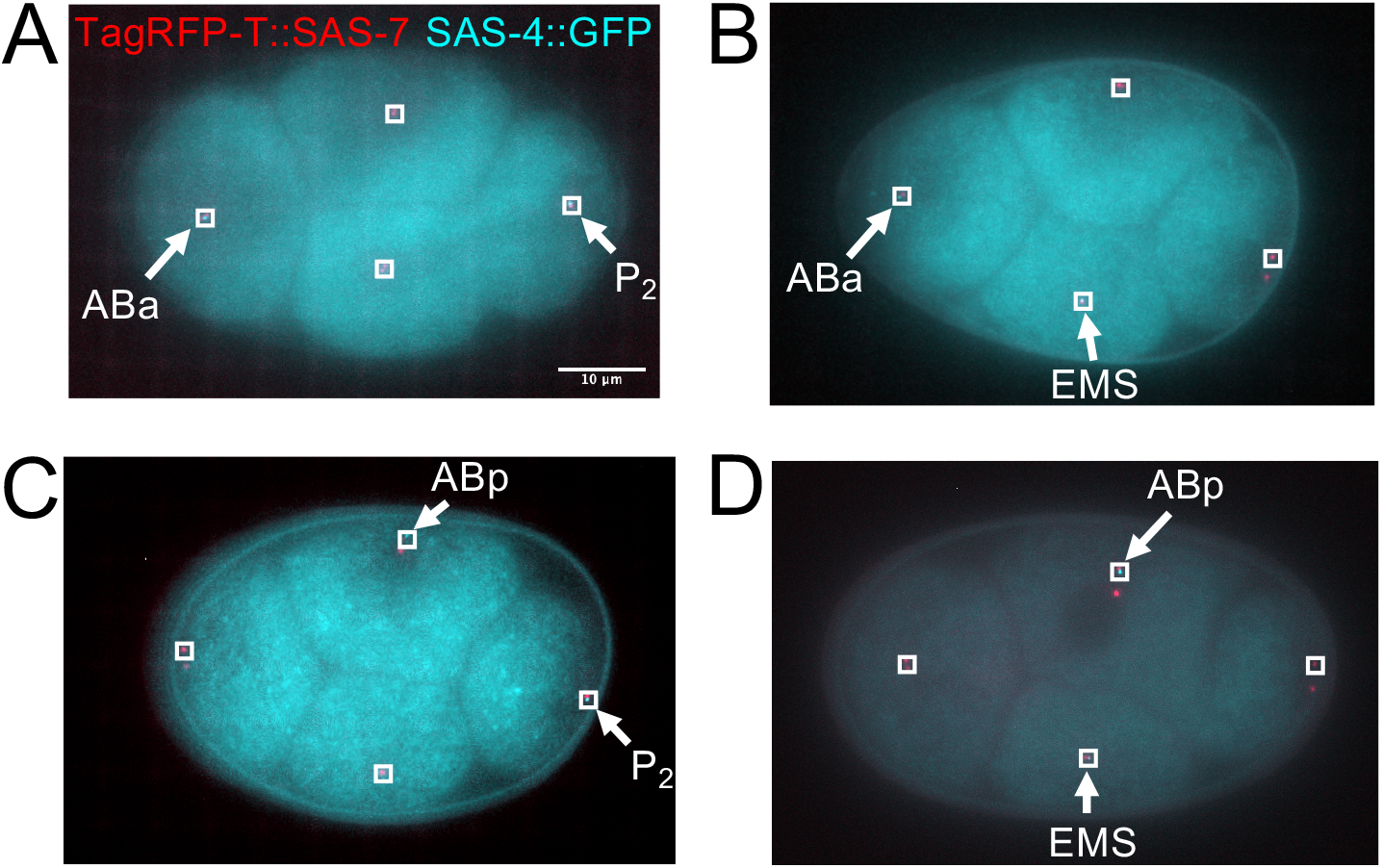
Four classes of sperm-contributed centriole distributions at the 4-cell stage. **(A-D)** Merged TagRFP-T::SAS-7 (red) and SAS-4::GFP (cyan) live imaging of 4-cell stage embryo resulting from the mating of TagRFP-T::SAS-7 *fem-1*(*hc17ts*) hermaphrodites with males expressing SAS-4::GFP. In each embryo, arrows point to the two blastomeres that inherited the sperm-contributed centrioles and insets correspond to the magnified views in Figure 2B-E (A: ABa and P_2_, same embryo as in Fig. 2A, B: ABa and EMS; C: ABp and P_2_; D: ABp and EMS).

**Video S1. Sequence of events in embryos derived from fertilization by sperm harboring a single centriole.**

DIC time-lapse recording of embryo derived from mating TagRFP-T::SAS-7 *fem-1*(*hc17ts*) hermaphrodites with *zyg-1*(*b1ts*) mutant males expressing SAS-4::GFP (not shown) and shifted during spermatogenesis to 25°C, yielding sperm with only one, older, centriole. Elapsed time since the beginning of the recording is shown in min:sec. The embryo is ~50 μm-long; anterior is to the left.

## Materials and Methods

### *C. elegans* strains

Worms were maintained using standard protocols (Brenner 1974). Strains of the following genotypes were utilized: *sas-7*(*Is1[tagRFP-T::sas-7+loxP]*) III, *fem-1(hc17ts)* IV (Doniach and Hodgkin 1984; Klinkert *et al.* 2019); *sas-7*(*or1940[gfp::sas-7]*) III (Sugioka *et al.* 2017); *sas-4*(*bs195[sas-4::gfp]*) III (a kind gift of Jessica Feldman and Kevin O’Connell); *zyg-1*(*b1ts*) II, *sas-4*(*bs195[sas-4::gfp*] III (Wood *et al.* 1980). Strains carrying temperature-sensitive alleles were maintained at 15°C, other strains at 24°C. To obtain hermaphrodites lacking sperm, *sas-7(Is1[tagRFP-T::sas-7+loxP]), fem-1(hc17ts)* were allowed to lay embryos overnight at 15°C on a plate, which was then shifted to 25°C for ~36-48 hours. The resulting sperm-less hermaphrodites were mated to *sas-4*(*bs195[sas-4::gfp]*) or *sas-7*(*or1940[gfp::sas-7]*) males, and their progeny analyzed by live imaging or following fixation (see below). To obtain sperm cells with a single centriole marked with SAS-4::GFP, *zyg-1*(*b1ts*), *sas-4*(*bs195[sas-4::gfp]*) males were grown initially at 15°C and shifted as L3 larvae to 25°C for ~24-48 hours prior to mating.

### Live imaging and analysis

Embryos were dissected from the uterus of hermaphrodites in a watch glass containing M9 and transferred using a mouth pipette onto a 2% agarose pad, which was overlaid gently with a 18×18 mm coverslip. Time-lapse DIC and dual color fluorescent microscopy imaging was performed at ~21°C on a Zeiss Axioplan 2 with a 63x 1.40 NA lens, with binning 2 and a 6% neutral density filter to attenuate the 120W Arc Mercury epifluorescent source. The motorized filter wheel, external shutters and the 1392 x 1040 pixels 12-bit Photometrics CoolSNAP ES2 camera were controlled by μManager (www.micro-manager.org). Typically, a z-stack of 15 planes 1 μm apart was taken at the 4-cell stage, with exposure times of 50 ms (DIC), 100 ms (GFP) and 100 ms (TagRFP-T). For display (Fig. 2A-E, Fig. S1), planes lacking centriolar signal were removed, and the remaining planes Max-intensity projected in Fiji. Brightness and contrast were adjusted slightly for better visualization, using identical settings within a series.

### Indirect immunofluorescence and confocal microscopy

Methanol fixation was performed essentially as described (Gönczy *et al.* 1999). In brief, gravid hermaphrodites were dissected on poly-lysine coated slides, covered with a 12×12 mm coverslip, and the slide frozen on a metal block pre-cooled on dry-ice. After a few minutes, embryos were freeze-cracked and fixed in −20°C methanol for ~3 min. Following two PBS washes, slides were incubated 45-60 min at room temperature with primary antibodies in PBT (PBS + 0.05% Tween-20). Primary antibodies were 1:50 mouse anti-IFA (Leung *et al.* 1999) and 1:500 rabbit anti-GFP (a kind gift of Viesturs Simanis). After two PBS washes, slides were incubated 45-60 min at room temperature with secondary antibodies in PBT (1:500 goat anti-rabbit-Alexa488 and goat anti-mouse-Alexa568 (ThermoFisher Scientific). Slides were counterstained with 1 μg/ml Hoechst 33258 (Sigma) to reveal DNA and washed twice with PBS. Thereafter, 7 μl of mounting medium (0.189 mol/l n-Propyl-Gallate, 90 % glycerol, 10 % PBS) was pipetted onto the specimen, which was covered with a 18×18 mm coverslip, applying slight pressure to remove excess liquid prior to analysis by wide-field microscopy. Indirect immunofluorescence was imaged on an upright LSM700 Zeiss confocal microscope with a Plan-Apochromat 63× 1.4 NA lens, collecting 1024×1024 pixels optical slices 0.35 μm apart, using 405 nm, 488 nm and 555 nm solid state lasers. Relevant planes containing centriolar signals were Max-intensity projected in Fiji. Brightness and contrast were adjusted slightly for better visualization, using identical settings within a series.

### Ethanol fixation

Gravid hermaphrodites were collected from plates with ~5 ml of M9, spun at 1500 rpm for 2 min in a clinical centrifuge, followed by two washes with M9. The supernatant was removed and 1.5 ml of 100% ethanol then added. Approximately 1 min thereafter, ethanol was removed, and the worm pellet resuspended in 15 μL MVD (50% M9, 50% Vectashield (Vector), 0.7 μg/L Hoechst 33258). After rehydration for 1 min or more, worms were transferred onto a slide and covered with a 20×40 mm coverslip, applying slight pressure to remove excess liquid prior to analysis of fixed embryos inside the uterus.

### Statistics

The presence of sperm-contributed centrioles in a given blastomere was determined and corresponding contingency tables generated using the Python library pandas. We then tested whether sister cells (ABa versus ABp, EMS versus P_2_), or any of the four possible distributions at the 4-cell stage (ABa and P_2_; ABa and EMS; ABp and P_2_; ABp and EMS), were more likely to harbor sperm-contributed centrioles using the χ^2^ test as implemented in the library SciPy ([https://scipy.org/citing-scipy/]). An analogous analysis was conducted with embryos that inherited a single sperm-contributed centriole. See Table S1 for details.

## Data Availability Statement

The authors affirm that all the data necessary for confirming the conclusions of the article are present within the article, figures, and tables. Strains are available upon request.

## Author contributions

Design: PG and FRB

Experiments: PG

Analysis: PG

Writing: PG

## Acknowledgements

We are grateful to Jessica Feldman and Kevin O’Connell for the SAS-4::GFP strain, as well as to Viesturs Simanis for GFP antibodies. Worm strains were provided also by the *Caenorhabditis* Genetics Center (CGC), which is funded by the NIH Office of Research Infrastructure Programs (P40 OD010440). We are grateful to Nils Kalbfuss and Alex Woglar for fruitful discussions and help in improving the manuscript, to Marie Pierron for advice, as well as to Léo Bürgy for help with statistical analysis.

---

**Table S1.**

| # sperm-contributed centrioles | figure | marker | comparison | n (embryos) | chisquare | p value |
| --- | --- | --- | --- | --- | --- | --- |
| 2 | Fig. 2F | GFP::SAS-4 | ABa vs ABp | 257 | 0.097 | 0.755 |
| 2 | Fig. 2F | GFP::SAS-4 | EMS vs P2 | 257 | 0.191 | 0.662 |
| 2 | Fig. 2G | GFP::SAS-4 | All equal? | 257 | 0.292 | 0.962 |
| 2 | Fig. 2J | GFP::SAS-7 | ABa vs ABp | 53 | 0.472 | 0.492 |
| 2 | Fig. 2J | GFP::SAS-7 | EMS vs P2 | 53 | 0.019 | 0.891 |
| 2 | Fig. 2K | GFP::SAS-7 | All equal? | 53 | 0.962 | 0.81 |
| 1 | Fig. 3D | GFP::SAS-4 | All equal? | 30 | 2.267 | 0.519 |

## References

Balestra F. R., L. von Tobel, and P. Gönczy, 2015 Paternally contributed centrioles exhibit exceptional persistence in *C. elegans* embryos. Cell research 25: 642–4. https://doi.org/10.1038/cr.2015.49

Blachon S., X. Cai, K. A. Roberts, K. Yang, A. Polyanovsky, et al., 2009 A proximal centriole-like structure is present in Drosophila spermatids and can serve as a model to study centriole duplication. Genetics 182: 133–44. https://doi.org/10.1534/genetics.109.101709

Bornens M., 2012 The centrosome in cells and organisms. Science 335: 422–426. https://doi.org/10.1126/science.1209037

Brenner S., 1974 The genetics of *Caenorhabditis elegans*. Genetics 77: 71–94.

Conduit P. T., and J. W. Raff, 2010 Cnn dynamics drive centrosome size asymmetry to ensure daughter centriole retention in Drosophila neuroblasts. Current biology: CB 20: 2187–92. https://doi.org/10.1016/j.cub.2010.11.055

Doniach T., and J. Hodgkin, 1984 A sex-determining gene, fem-1, required for both male and hermaphrodite development in *Caenorhabditis elegans*. Dev Biol 106: 223–235. https://doi.org/10.1016/0012-1606(84)90077-0

Erpf A. C., and T. Mikeladze-Dvali, 2020 Tracking of centriole inheritance in *C. elegans*. MicroPubl Biol 2020. https://doi.org/10.17912/micropub.biology.000256

Fawcett D. W., 1975 The mammalian spermatozoon. Dev Biol 44: 394–436. https://doi.org/10.1016/0012-1606(75)90411-x

Gönczy P., H. Schnabel, T. Kaletta, A. D. Amores, T. Hyman, et al., 1999 Dissection of cell division processes in the one cell stage *Caenorhabditis elegans* embryo by mutational analysis. The Journal of cell biology 144: 927–946.

Januschke J., S. Llamazares, J. Reina, and C. Gonzalez, 2011 Drosophila neuroblasts retain the daughter centrosome. Nature communications 2: 243. https://doi.org/10.1038/ncomms1245

Kirkham M., T. Müller-Reichert, K. Oegema, S. Grill, and A. A. Hyman, 2003 SAS-4 is a *C. elegans* centriolar protein that controls centrosome size. Cell 112: 575–587.

Klinkert K., N. Levernier, P. Gross, C. Gentili, L. von Tobel, et al., 2019 Aurora A depletion reveals centrosome-independent polarization mechanism in *C. elegans*. bioRxiv. https://doi.org/10.1101/388918

Leidel S., and P. Gönczy, 2003 SAS-4 is essential for centrosome duplication in *C elegans* and is recruited to daughter centrioles once per cell cycle. Developmental cell 4: 431–439.

Leung B., G. J. Hermann, and J. R. Priess, 1999 Organogenesis of the *Caenorhabditis elegans* intestine. Developmental biology 216: 114–34.

Nigg E. A., and T. Stearns, 2011 The centrosome cycle: Centriole biogenesis, duplication and inherent asymmetries. Nature cell biology 13: 1154–1160. https://doi.org/10.1038/ncb2345

O’Connell K. F., C. Caron, K. R. Kopish, D. D. Hurd, K. J. Kemphues, et al., 2001 The *C. elegans zyg-1* gene encodes a regulator of centrosome duplication with distinct maternal and paternal roles in the embryo. Cell 105: 547–558.

Sugioka K., D. R. Hamill, J. B. Lowry, M. E. McNeely, M. Enrick, et al., 2017 Centriolar SAS-7 acts upstream of SPD-2 to regulate centriole assembly and pericentriolar material formation. eLife 6. https://doi.org/10.7554/eLife.20353

Wang X., J. W. Tsai, J. H. Imai, W. N. Lian, R. B. Vallee, et al., 2009 Asymmetric centrosome inheritance maintains neural progenitors in the neocortex. Nature 461: 947–55. https://doi.org/nature08435 [pii] 10.1038/nature08435

Winey M., and E. O’Toole, 2014 Centriole structure. Philos Trans R Soc Lond B Biol Sci 369. https://doi.org/10.1098/rstb.2013.0457

Wolf N., D. Hirsh, and J. R. McIntosh, 1978 Spermatogenesis in males of the free-living nematode, *Caenorhabditis elegans*. Journal of ultrastructure research 63: 155–169.

Wood W. B., R. Hecht, S. Carr, R. Vanderslice, N. Wolf, et al., 1980 Parental effects and phenotypic characterization of mutations that affect early development in *Caenorhabditis elegans*. Dev. Biol. 74: 446–469.

Yamashita Y. M., A. P. Mahowald, J. R. Perlin, and M. T. Fuller, 2007 Asymmetric inheritance of mother versus daughter centrosome in stem cell division. Science 315: 518–521.

Yamashita Y. M., 2009 Regulation of asymmetric stem cell division: spindle orientation and the centrosome. Frontiers in bioscience: a journal and virtual library 14: 3003–11.

